# An optimized model for HEV infection in the HepaRG cell line

**DOI:** 10.1101/2025.08.01.667596

**Authors:** Stacy Gellenoncourt, Marie Pellerin, Aïlona Marcadet-Hauss, Roxanne Fouillé, Michel Rivoire, Guillaume Passot, Julie Lucifora, David Durantel, Nicole Pavio, Virginie Doceul

**Affiliations:** Institut National de Recherche pour l’Agriculture, l’Alimentation et l’Environnement (INRAE), Agence Nationale de Sécurité Sanitaire de l’Alimentation, de l’Environnement et du Travail (ANSES), École Nationale Vétérinaire d’Alfort (ENVA), UMR Virologie, Maisons-Alfort, France; Centre International de Recherche en Infectiologie (CIRI), Univ Lyon, Inserm, U1111, Université Claude Bernard Lyon 1, CNRS, UMR5308, ENS de Lyon, F-69007, Lyon, France; Centre Léon Bérard (CLB), INSERM, U1032, Lyon, France; Service de chirurgie générale et oncologique, Hôpital Lyon Sud, Hospices Civils de Lyon Et CICLY, EA3738, université Lyon 1

**Keywords:** Hepatitis E virus, HepaRG, DMSO, Differentiation, Innate immunity

## Abstract

Hepatitis E virus (HEV) causes acute hepatitis that can progress to fulminant or chronic hepatitis. For decades, the lack of a pertinent and robust cell culture system for HEV has delayed our understanding on this hepatotropic virus. HepaRG cells are one of the few hepatocyte-derived cell lines able to replicate HEV. These cells can differentiate (dHepaRG) into hepatocytes and cholangiocytes upon treatment with dimethyl sulfoxyde (DMSO) and are very relevant to study interactions between pathogens and hepatocyte innate immunity. However, the suitability of the HepaRG model to study HEV need to be further investigated. In this study, we found that HEV can infect proliferating HepaRG cells and that DMSO-induced differentiation is not necessary for HEV infection. Moreover, even if treatment with DMSO is needed to maintain optimal differentiation and polarization of dHepaRG, its presence is detrimental for HEV infection. Overall, this study shows that dHepaRG cells cultured without DMSO is a suitable model to study HEV and its interaction with the hepatocyte innate system.

## 1. Introduction

Hepatitis E virus (HEV, *Paslahepevirus balayani*) is responsible for acute hepatitis in humans. Infections are generally asymptomatic but can evolve to fulminant hepatitis, particularly in pregnant women, or cause chronic hepatitis in immunocompromised patients (Li et al., 2020). WHO estimates that around 20 million of acute infections occur per year (“Hepatitis E,” n.d.). HEV is spread worldwide, with an average seroprevalence of 12.5% and an incidence that has increased by 18% between 1990 and 2017 (Jing et al., 2021). A rise in the number of cases has also been described from 2010 to 2019 but might be due to improved diagnosis and surveillance of asymptomatic and mild courses (Schemmerer et al., 2022). Eight HEV genotypes have been identified (named HEV-1 to -8) and differ by their host range. HEV-1 and -2 are restricted to humans while HEV-3, -4 and -7 are zoonotic (Kinast et al., 2022; Lee et al., 2016). Recently, HEV has been classified in the top ten of viruses presenting a high risk of zoonotic spill over from wild fauna among 800 studied viruses (Grange et al., 2021). To improve our knowledge on HEV, robust and pertinent *in vitro* models are needed (reviewed in (Fu et al., 2019)). However, development of such *in vitro* models has proven to be a real challenge due to low infection level and slow replication in multiple cell lines. Primary human hepatocytes (PHH) are the closest cellular model to human liver that can be used to study hepatitis viruses as well as metabolism or compound toxicity (Gupta et al., 2021; Schulze et al., 2012; Xiao et al., 2021). PHH transcriptome is closer to the one from liver samples than commonly used hepatic cell lines such as HepaRG or HepG2 (Gupta et al., 2021). However, their accessibility and their maintenance *in vitro* are limited (Sugahara et al., 2023; Weaver et al., 2023; Xiao et al., 2021). Moreover, studies using PHH have to take into account inter-individual variability (Jetten et al., 2013; Parmentier et al., 2018) even if improvements have been made to overcome sourcing, divergence (Michailidis et al., 2020; Uehara et al., 2023) and viability (Weaver et al., 2023; Xiao et al., 2021) issues. To bypass those limitations, different cell lines have been developed including the HepaRG cell line. HepaRG cells are bipotent progenitor cells able to differentiate into functional hepatocytes and biliary-like epithelial cells (Parent et al., 2004). Differentiation of HepaRG cells is usually reached after addition of dimethyl sulfoxyde (DMSO) in the culture media (Gripon et al., 2002). DMSO exerts epigenetic alterations, mainly through histone modifications, leading to cell differentiation by promoting the expression of liver-specific genes implicated in metabolism or xenobiotic detoxification associated to the liver functions (Dubois-Pot-Schneider et al., 2022; Sugahara et al., 2023). HepaRG cells are a good alternative to PHH as they can be used for studies on toxicity, metabolism (Andreozzi et al., 2023; Yokoyama et al., 2018) and virus/host interactions (Gripon et al., 2002; Luangsay et al., 2015; Zhang et al., 2018). It has been showed that HepaRG cells are closer to PHH than HepG2 cells, regarding expression of genes involved in liver-related pathways (Ardisasmita et al., 2022; Stanley and Wolf, 2022). Moreover, expression of sensors of innate immunity in HepaRG cells is closer to PHH than other hepatocyte cell lines and HepaRG cells are functional for IFN signalling (Luangsay et al., 2015; Lucifora et al., 2018). HepaRG cells have been identified as a robust model to study Hepatitis B virus (HBV) (Gripon et al., 2002; Li et al., 2016; Yan et al., 2013) and Hepatitis D virus (HDV) (Alfaiate et al., 2016; Li et al., 2016; Yan et al., 2013, 2012). DMSO-induced differentiation of HepaRG increases expression of the HBV entry receptor, sodium taurocholate cotransporting polypeptide (NTCP) (Gripon et al., 2002; Yan et al., 2012), and is mandatory to ensure HBV entry. In contrast to the well characterized HepaRG model for HBV infection, only few studies have shown that HEV can replicate in HepaRG cells (Klöhn et al., 2024; Pellerin et al., 2021; Rogée et al., 2013; Schrader et al., 2023) and none has assessed the importance of HepaRG differentiation state on HEV infection.

In this study, we assessed the impact of DMSO-induced differentiation on HEV infection in HepaRG cells and further evaluated the pertinence of this model to study HEV. We found that DMSO-induced differentiation of HepaRG cells is not necessary for optimal HEV infection and that maintenance of DMSO during infection has a negative impact on HEV-3 infection despite the maintenance of a better differentiation and polarization profile. Overall, this study provides new evidence of the usefulness and suitability of the HepaRG cell model to study HEV-3 infection and its interaction with the hepatocyte innate system.

## 2. Materials and methods

### 2.1. Cell culture

Undifferentiated human HepaRG^TM^ cells were purchased from BIOPREDIC International. Cells were grown in “proliferation medium” consisting of William’s E medium with GlutaMAX^TM^ (ThermoFisher Scientific) supplemented with 10% heat-inactivated fetal bovine serum (FBS, EuroBio), 5 μg/ml insulin (Sigma-Aldrich, #I9278), 5×10^−5^ M hydrocortisone hemisuccinate (Sigma-Aldrich, #1319002) and 100 IU/ml penicillin and 100 µg/ml streptomycin. Cells were maintained at 37°C in 5% CO_2_. Confluent HepaRG monolayers were trypsinated every 2 weeks and medium was renewed twice a week. For differentiation, HepaRG were seeded into 6-or 24-well plate and cultured in proliferation medium for 2 weeks corresponding to day 1 (D1) post-plating (p.p) to D14p.p. Medium was then replaced for 14 days (D14p.p to D28p.p) by “differentiation medium” consisting of HepaRG proliferation medium supplemented with 1.2% DMSO (Sigma-Aldrich, #D2650). HepaRG cells that underwent this 2-week treatment with DMSO are referred as differentiated HepaRG (dHepaRG). HepaRG cells were then maintained in differentiation medium (+ DMSO) or proliferation medium (-DMSO) accordingly. Primary human hepatocytes (PHH) were isolated from human liver resections obtained from the Centre Léon Bérard (Lyon) and Hopital de Lyon Sud with French ministerial authorizations (AC 2013-1871, DC 2013 – 1870, AFNOR NF 96 900 sept 2011) as previously described (LeCluyse and Alexandre, 2010). Informed consent was obtained from all subjects and/or their legal guardian(s). PHH were seeded and cultured for 24h before lysis for further analyses.

### 2.2. HEV-3 infection of HepaRG

A HEV-3f strain originating from a French patient suffering from acute autochthonous hepatitis E was used and has been described previously (GenBank under accession number JN906974) (Bouquet et al., 2012; Pellerin et al., 2021). Supernatant from the 7th passage of the virus in HepaRG cells was used in this study to perform infections in HepaRG cells. Cells were counted prior to infection (at D3, D7, D14 and D30p.p) and infected at the specified MOI (indicated in genome equivalent (GE)/cell). Cells were then incubated overnight with the HEV inoculum in a final volume of 1 mL, 500 µL or 250 µL for 6-, 12- and 24-well plate respectively, before being washed three times with PBS the next day. Cells were maintained in proliferation medium (-DMSO) or differentiation medium (+DMSO) until the end of the experiment as indicated and the medium changed 2 to 3 times per week.

### 2.3. Viral RNA extraction from supernatant

Viral RNA was extracted from 200μl culture supernatants using the MagMAX core nucleic acid purification kit (Thermo Fisher Scientific, #A32702) and the KingFisher instrument according to the manufacturer instructions and as described previously (Pellerin et al., 2022).

### 2.4. Intracellular RNA extraction

HepaRG cells were washed 3 times in cold PBS and harvested. Total RNA was extracted using the RNeasy minikit (Qiagen, #74106) including a digestion step on column with DNase I (Qiagen, #79254) according to the supplier’s protocol. After a first elution a second digestion step was performed with TurboDNase (Invitrogen, #10646175) according to the supplier’s protocol. After digestion, purification was done based on the RNA cleaned up procedure from the RNeasy minikit (Qiagen). RNA concentrations were determined and kept at -80°C until use. Total intracellular RNA from PHH were extracted with the “Monarch, nucleic acid purification kit” according to the manufacturer’s instructions (New England Biolabs). The use of primary human hepatocytes was approved by French ministerial authorizations (AC 2013-1871, DC 2013 – 1870, AFNOR NF 96 900 sept 2011) and conducted in accordance with the local legislation and institutional requirements.

### 2.5. Quantification of cellular gene expression by RT-qPCR

Reverse transcription was performed using 500 ng of RNA with SuperScript IV Reverse Transcriptase (Fisher, #15307696). RT-qPCR was performed on 2 µl of cDNA (pre-diluted by half in DNAse-free water) using the QuantiTect SYBR Green PCR Kit (Qiagen, #204145), and specific primers as listed (Table 1). A LightCycler 480 apparatus (Roche) was used for sample analysis. Samples were denatured for 15 min at 95°C, then DNA was amplified with 40 cycles at 95°C for 30 s, 60°C for 30 s. The final extension was followed by cooling at 40°C for 30 s. *GAPDH*, *B2M* and *GUSB* were used as housekeeping genes for normalization and cells harvested at D4p.p were used as reference condition. Relative gene expression was determined using the Vandesompele method (Hellemans et al., 2007).

**Table 1:**
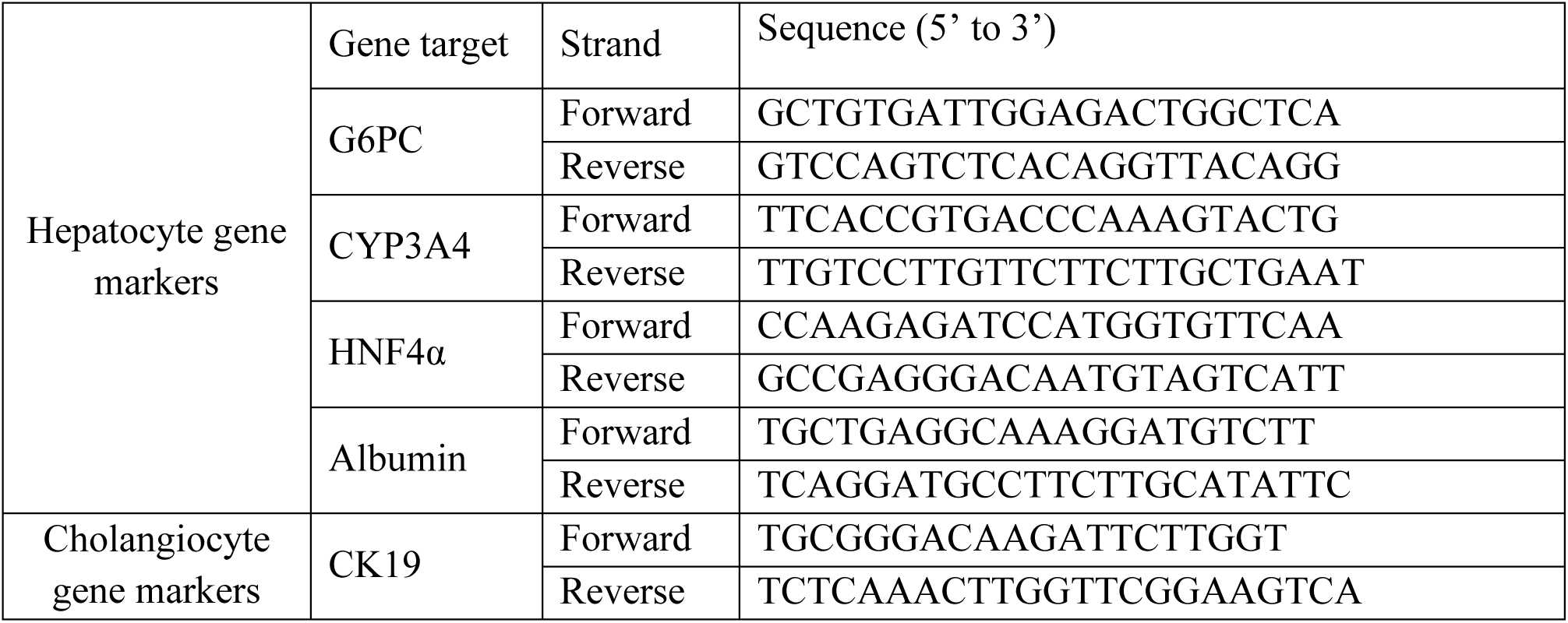
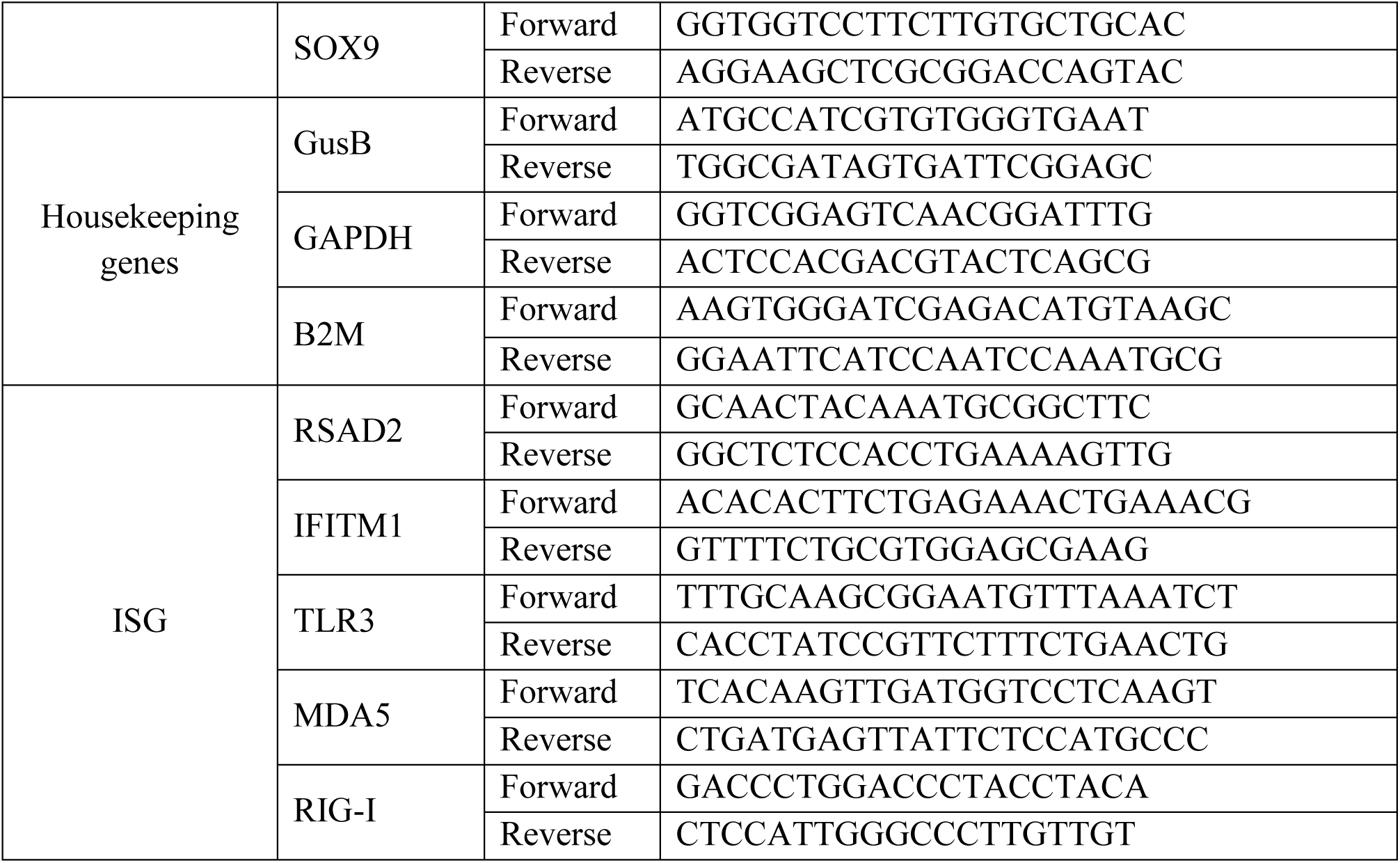
List of primers used for the relative quantification of human gene expression by RT-qPCR.

### 2.6. HEV quantification by real-time quantitative PCR (RT-qPCR)

HEV RNA quantification was adapted from the method described by Jothikumar et al. (2006) as described previously (Barnaud et al., 2012). The QuantiTect Probe RT-PCR kit (Qiagen) was used according to the manufacturer’s instructions using 2 μl of RNA (template), 0.25 mM reverse primer (5’-AGGGGTTGGTTGGATGAA-3’), 0.1 mM forward primer (5’-GGTGGTTTCTGGGGTGAC-3’) and 5mM probe (FAMTGATTCTCAGCCCTTCGC-MGB). A LightCycler 480 apparatus (Roche Molecular Biochemicals) and its associated software (LightCycler 480 Software release, v.1.5.039) was used for sample analysis. Reverse transcription was carried out at 50°C for 20 min, followed by denaturation at 95°C for 15 min. DNA was amplified with 45 cycles at 95°C for 10 s and 58°C for 45 s. Standard HEV RNA was obtained after *in vitro* transcription of a plasmid pCDNA 3.1 ORF2-3 HEV and used to generate standard quantification curves as described previously (Barnaud et al., 2012).

### 2.7. Immunostaining and fluorescent microscopy

Cells were seeded onto a 15µ-24-well IBIDI plate (Clinisciences). After infection, cells were washed in PBS and fixed with 4% paraformaldehyde in PBS. Cells were then permeabilised with 0.2% Triton X-100 in PBS for 1h before being incubated in blocking buffer (0.5% BSA) for 30 min. A polyclonal anti-ORF2 antibody from rabbit sera (gift from Katja Dinkelborg (Ssebyatika et al., 2025)), mouse anti-MRP2 (1:50, Cliniscience #MON9027C) and/or goat anti-albumin (1:500, Thermofisher #A80129A) were then incubated overnight at 4°C in PBS containing 5% BSA. Cells were then washed several times in PBS and incubated with the appropriate secondary antibody: AF488 donkey anti-mouse (1:500, InVitrogen #A32766), AF546 donkey anti-goat (1:500, Thermofisher #10153892), AF488 donkey anti-rabbit (1:500, Thermofisher #10424752). After several washes in PBS, cell nuclei were stained with 4,6-diamidine-2-phenylindole dihydrochloride (DAPI) (Sigma-Aldrich) for 10 min. Images were acquired using an Axio observer Z1 fluorescent microscope (Zeiss) and the Zen 2012 software (version 8,0,0,273).

### 2.8. IFN-β1 stimulation

HepaRG were cultivated and differentiated as mentioned before and kept in proliferation medium supplemented or not with DMSO until the end of the experiment. Cells were stimulated with 200UI/mL of IFN-β1 (PBL Interferon Source, Piscataway, NJ, USA) overnight in presence or in absence of DMSO at D30 and D57p.p. Cell pellets were harvested the next day after 3 washes with PBS and kept at -80°C until use.

### 2.9. Flow Cytometry

At specific time points post-infection, HepaRG cells were harvested and frozen in FBS containing 10% DMSO. Cells were then thawed and washed in PBS before being stained for viability (1:400, Thermofisher # 65-0865-14) for 20 min at 4°C. Cells were washed twice in PBS before being fixed and permeabilized with methanol at -20°C for 1h. Methanol was removed and cells were washed with PBS. Cells were then stained with mouse anti-ORF2 (1:200, Milipore #MAB8002) diluted in Perm Wash buffer (BD #562574) for 45 min at 37°C. Cells were washed twice in PBS and stained with secondary antibody, donkey anti-mouse 488 (1:200, Thermofisher #10153892) for 90 min at 4°C. Finally, cells were washed in PBS before acquisition on BD FACS Canto2. Flowjo software was used to analyse data (version 10.8.1).

### 2.10. Statistical analysis

Statistical tests are indicated in each legend figure and were done on GraphPad Prism 9 (version 9.2.0).

## 3. Results

### 3.1. DMSO-induced differentiation is not essential for HEV infection in HepaRG cells

To differentiate HepaRG cells into hepatocytes and cholangiocytes, cells are seeded for 2 weeks into plates (corresponding to the proliferation phase) before addition of 1-2 % DMSO for 2 additional weeks to the culture media (Gripon et al., 2002). DMSO is then kept in the culture media to maintain differentiation (Dubois-Pot-Schneider et al., 2022). Morphologically, HepaRG cells first proliferate to reach confluence 1 to 2 weeks after plating. DMSO is then added for 2 extra weeks and the formation of hepatocyte-like cells as well as cholangiocyte-like cells is visible (Fig. 1A). To assess whether DMSO-induced differentiation is essential for HEV infection, HepaRG cells were infected with HEV-3 at a MOI of 10 genome equivalent (GE) / cell at different stages of cell proliferation without any addition of DMSO: at 3 (Fig. 1B, green), 7 (Fig. 1B, black), 14 (Fig. 1B, orange) and 30 (Fig. 1B, purple) days post-plating (p.p.).

**Figure 1:**
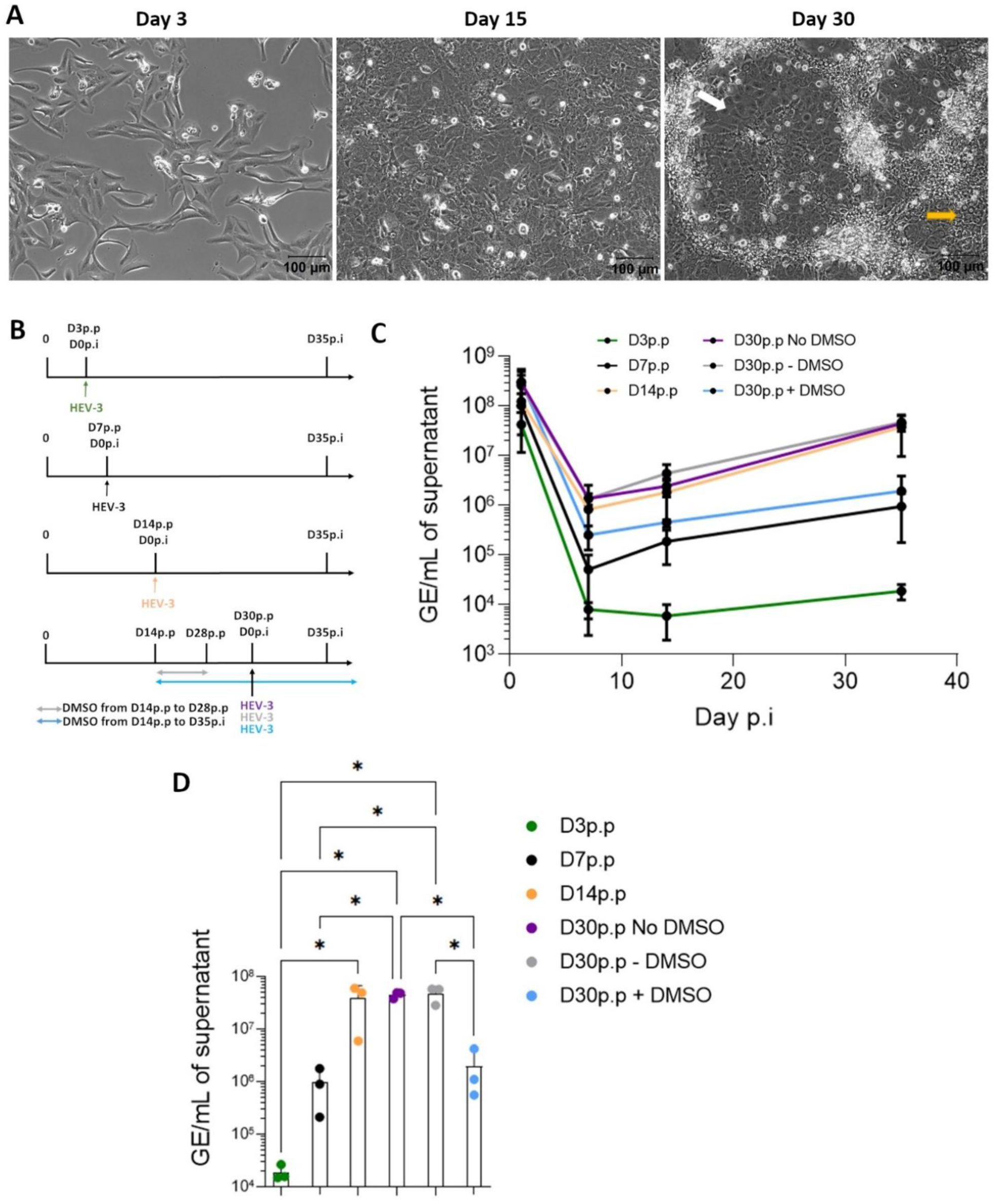
HEV-3 can infect proliferating and differentiated HepaRG cells. **(A)** Images of HepaRG cells at different stages of proliferation (D3p.p and D15p.p) and after DMSO differentiation (D30p.p). White and orange arrows show examples of cholangiocyte-like and hepatocyte-like cells, respectively. Scale bar is 100µm. **(B)** Experimental design of infection of HepaRG at MOI 10 at day 3 (green), 7 (black) and 14p.p (orange) or D30p.p in absence of DMSO (purple), with DMSO from D14 until D28p.p (grey) or with DMSO from D14p.p until D35p.i (blue). **(C)** Mean viral load in supernatant (GE/mL, ±SD) over time after infection under different conditions as described in (B). Each independent experiment (n≥2) was performed in triplicates. **(D)** Mean viral load (±SD) in supernatant at D35p.i as represented in (C). One-way ANOVA with Tukey’s multiple comparisons, *: *p-value*<0.05. p.p: post-plating, p.i: post-infection, MOI 10: 10 genome equivalent (GE)/cell.

In parallel, differentiated HepaRG (dHepaRG) cells that were cultivated for 2 weeks in DMSO (D14 to D28 p.p.) were also infected at D30p.p in the absence (Fig. 1B, grey) or presence of DMSO (Fig. 1B, blue). Viral RNA loads were then quantified in the cell supernatant over time (Fig. 1C). After infection at 3 or 7 days p.p, increased level of HEV-3 RNA was measured (Fig. 1C, green and black), showing that HEV-3 can infect proliferating HepaRG cells. However, higher level of HEV RNA was obtained after infection of confluent HepaRG cells at D14p.p or D30p.p (Fig. 1C-D, orange and purple). Interestingly, the addition of DMSO for 2 weeks prior to infection at D30p.p resulted in a similar kinetic of HEV-3 RNA secretion (Fig. 1C-D, grey), suggesting that DMSO-induced differentiation does not enhance HEV-3 infection. Moreover, maintenance of DMSO during infection in dHepaRG had a negative impact on HEV-3 infection (Fig. 1C-D, blue) as compared to dHepaRG maintained without DMSO (Fig. 1C-D, grey). Overall, these results suggest that DMSO-induced differentiation is not essential for HEV-3 infection and that presence of DMSO during infection has a detrimental effect on HEV-3 infection.

### 3.2. HEV RNA levels are reduced in dHepaRG cells cultivated with DMSO during infection

Next, to confirm that HEV-3 replication was higher in dHepaRG maintained without DMSO, dHepaRG cells were infected (MOI 100 GE/cell) in the presence or absence of DMSO for longer period of time (Fig. 2A). As observed before, extracellular HEV-3 viral load was higher in absence of DMSO compared to the condition with DMSO. This difference was maintained until D100 post-infection (p.i) (Fig. 2B). Intracellular HEV-3 RNA was also measured at D100p.i and lower level of intracellular HEV-3 RNA was also detected in dHepaRG maintained in DMSO in comparison to the one maintained without DMSO (Fig. 2C). In accordance with this result, lower number of ORF2 positive cells was also detected by immunofluorescent staining at D100 p.i (Fig. 2D) in the presence of DMSO. Proportion of infected cells was evaluated by flow cytometry. Less than 5% of HEV-3 infected cells were detected in absence of DMSO at D50p.i. In dHepaRG maintained in DMSO, the percentage was lower with 0.2% of infected cells detected at D50p.i (Supplementary Fig. S1).

**Figure 2:**
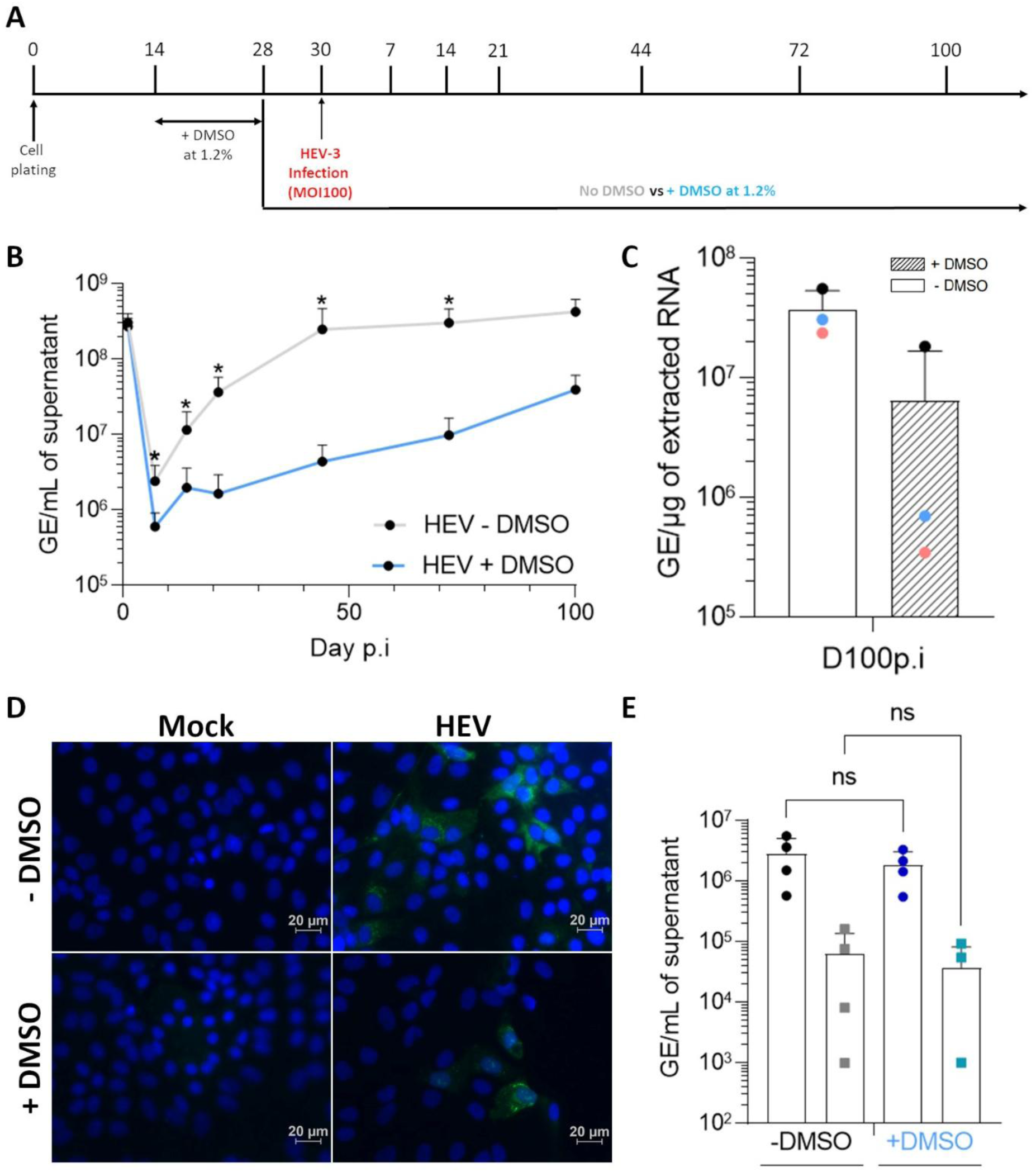
Optimal HEV-3 replication is detected in dHepaRG cultivated in absence of DMSO. **(A)** HepaRG cells were plated 2 weeks before addition of 1.2% of DMSO for another 2 weeks to obtain differentiated HepaRG cells (dHepaRG). At day 28 p.p, DMSO was kept (blue) or removed (grey) before HEV-3 infection (MOI 100) at D30p.p. Cells were maintained until day 100p.i (corresponding to D130p.p) in presence or in absence of DMSO. **(B)** Mean HEV-3 viral load (GE/mL, ±SD) in supernatant of infected dHepaRG overtime in presence (blue) or in absence (grey) of DMSO. Each independent experiment (n≥3) was performed in triplicates. **(C)** Mean intracellular viral load (GE/µg of extracted RNA, ±SD) at D100p.i in infected dHepaRG in presence (hatched) or in absence (white) of DMSO. Each independent experiment (n=3) was performed in triplicates. **(D)** Immunofluorescence microscopy of infected (MOI 100) dHepaRG at D100p.i in absence (top) or in presence of DMSO (down). Staining specific to nuclei (DAPI, blue) and HEV ORF2 (green) is shown. Scale bar is 20µm. **(E)** Mean viral load (GE/mL, ±SD) in supernatant at D35p.i from dHepaRG maintained without DMSO and infected with HEV-3 produced on dHepaRG maintained in presence (blue points) or in absence of DMSO (dark points) at different MOI (10: •, 1: □). Each independent experiment (n=4) was performed in duplicates. (B-C) Mann-Whitney, (E) ANOVA with Dunn’s multiple comparison *: *p-value*<0.05. p.p: post-plating, p.i: post-infection, MOI 100: 100 genome equivalent (GE)/cell.

### 3.3. DMSO does not impact infectivity of viral particles

The impact of DMSO on the infectivity of viral particles produced in dHepaRG cells was also assessed. To do this, supernatant from HEV-infected dHepaRG, cultivated in absence or in presence of DMSO, was used to infect newly-prepared dHepaRG at different MOI (10 and 1 GE/cell) under the same conditions. At D35p.i, the same viral load was detected in the supernatant of infected cells, independently of the inoculum used (Fig. 2E). This result suggests that maintenance of DMSO during HEV infection in HepaRG cells impairs the quantity of viral particles released in the supernatant but not their infectivity.

### 3.4. dHepaRG maintained without DMSO are in an intermediate differentiation and polarization state

Then, we assessed the differentiation state of HepaRG cells upon culture in presence or absence of DMSO by measuring the expression level of genes coding for different hepatocyte and cholangiocyte markers by RT-qPCR. Hepatocytes express high level of albumin (*ALB*) (Dianat et al., 2014; Gurevich et al., 2020), cytochrome P450 (*CYP3A4*) (Gurevich et al., 2020; Peng et al., 2018) as well as glucose-6-phosphatase (*G6PC*) (Bonanini et al., 2024; Yang et al., 2023). On the other hand, sex-determining region Y-Box transcription 9 (*SOX9*), cytokeratin 19 (*CK19)* (Cerec et al., 2007; Higuchi et al., 2014) are considered as cholangiocytes markers. The level of expression of these markers are commonly used to follow the differentiation of HepaRG (Dubois-Pot-Schneider et al., 2022; Higuchi et al., 2014; Mayati et al., 2018; Prabhakar et al., 2021).

Hepatocyte gene expression was measured in uninfected dHepaRG cells at different time points until D130p.p as well as in HEV-3 infected dHepaRG at D100p.i (corresponding to D130p.p) and in uninfected PHH. For all conditions, including PHH, level of expression of the genes coding for the different hepatocyte (Fig. 3A) and cholangiocyte (Fig. 3B) markers was normalized to the one of the HepaRG harvested 4 days post-plating. At confluency (D14p.p), the level of expression of the genes *G6PC*, *CYP3A4* and *ALB* was upregulated in HepaRG, showing that, even before DMSO addition, cells already started to differentiate. After 2 weeks of culture (D28p.p) in the presence of DMSO, gene expression of these hepatocyte markers increased significantly and was similar or higher in comparison to PHH, confirming the suitability of the HepaRG cell line as an alternative to PHH. However, after removal of the DMSO for only 2 days (D30p.p), expression of the hepatocyte markers decreased and was lower in comparison to the cells maintained in DMSO. This difference between cells maintained in presence or in absence of DMSO was maintained over time. Interestingly, gene expression remained higher than basal level (D4p.p) in the absence of DMSO until D130p.p, suggesting the persistence of an intermediate differentiation state (Fig. 3A). The level of expression of the cholangiocyte marker genes, *CK19* and *SOX9*, was slightly modulated by DMSO-induced differentiation in HepaRG (D30p.p) (Fig. 3B). A decrease in CK19 expression level was also observed in dHepaRG after long culture in the presence of DMSO (Fig. 3B).

**Figure 3:**
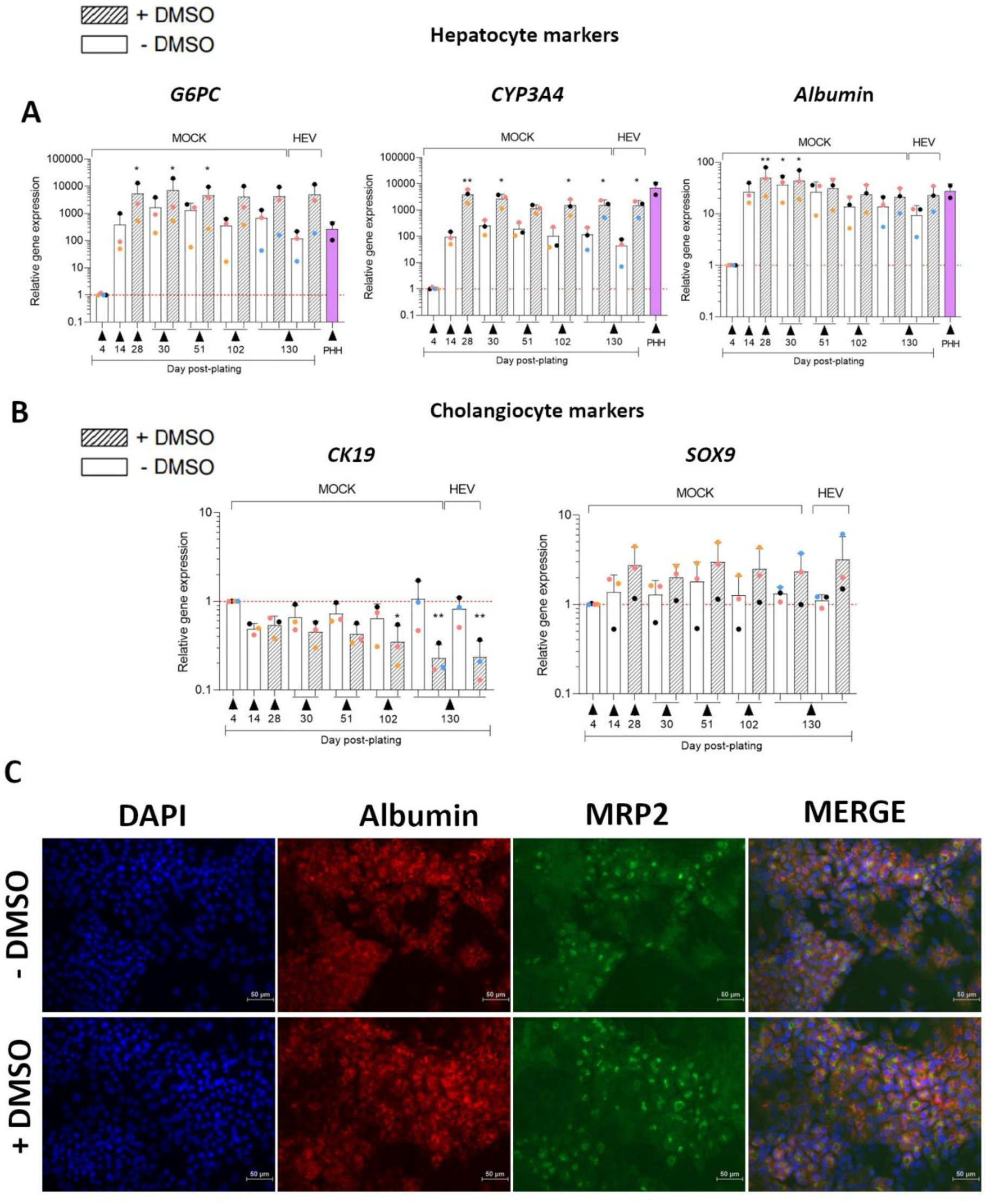
dHepaRG cells maintained without DMSO are in an intermediate state of differentiation. **(A, B)** Mean relative gene expression of hepatocyte (A) or cholangiocyte (B) marker genes in non-infected PHH or dHepaRG (MOCK) in presence (hatched) or in absence (white) of DMSO upon time. Relative gene expression in dHepaRG infected with HEV-3 (HEV) at MOI 100 in presence (hatched) or in absence (white) of DMSO is also shown at D130p.p. Relative gene expression (±SD) was calculated compared to D4p.p. *GusB, B2M* and *GAPDH* were used as housekeeping genes. (A-B) Kruskal-Wallis test compared to D4p.p where each independent experiment (n≥2) was performed in triplicates, * *p-value*<0.05, ** *p-value*<0.01. **(C)** Immunofluorescence microscopy on dHepaRG at D30p.p in absence (top) or in presence of DMSO (bottom). Staining specific to nuclei (DAPI, blue), MRP2 (green) and albumin (red) is shown. p.p: post-plating, PHH: primary human hepatocytes. Scale bar is 50µm.

The effect of DMSO on the protein level of ALB was also assessed by immunofluorescence (IF) microscopy. Staining of ALB was mostly observed in hepatocyte-like cells as previously reported (Schrader et al., 2023) (Fig. 3C). No major impact of DMSO removal (for 2 days and 23 days) was observed by IF microscopy at D30p.p (Fig. 3C) or at D51p.p (Supplementary Fig. S2A).

Polarization of cells allows directional trafficking and import/export of molecules/bile (Bachour-El Azzi et al., 2015; Le Vee et al., 2013). One of the transporters implicated in bile secretion is an organic anion transporter called multidrug resistance associated protein 2 (MRP2) (Higuchi et al., 2014) which expression can be induced by DMSO (Mayati et al., 2018). To address whether HepaRG cells remain polarized, we assessed MRP2 protein expression by IF microscopy 2 days after DMSO removal (D30p.p). MRP2 was detected in both conditions (Fig. 3C). However, we observed a loss of MRP2 expression upon prolonged culture in absence of DMSO (Supplementary Fig. S2B).

Overall, these results show that the presence of DMSO is essential to maintain optimal differentiation and polarization of HepaRG but that dHepaRG cells remain in an intermediate state after DMSO removal for prolonged time.

### 3.5. Presence of DMSO does not impact expression level of innate immune sensors

We then assessed whether dHepaRG cells cultivated without DMSO would be a pertinent model to study interaction between the host innate immunity and HEV. First, we evaluated the impact of DMSO on the expression level of *retinoic-acid-inducible gene I* (*RIG-I), melanoma differentiation-associated protein 5* (*MDA5*) and *Toll-like receptor 3* (*TLR3*) that have been shown to play a role in the sensing of HEV (Devhare et al., 2016; Xu et al., 2017; Yin et al., 2017). Expression levels of the genes coding for *RIG-I, MDA5* and *TLR3* were followed overtime in dHepaRG in the presence or absence of DMSO. Results showed no modulation of the expression of *RIG-I*, *MDA5* and *TLR3* gene expression by DMSO until D102p.p in uninfected dHepaRG cells. Moreover, gene expression levels seemed close to the one of fresh PHH (Fig. 4A). We also assessed the ability of dHepaRG to respond to IFN-I stimulation in the presence or absence of DMSO. Uninfected dHepaRG, maintained or not in DMSO, were incubated with IFN-β1 at D31 and D58p.p and the level of expression of 2 interferon-stimulated genes (ISGs), *radical S-adenosyl methionine domain containing 2* (*RSAD2)* and *interferon induced transmembrane protein 1 (IFITM1*), analysed 16h after stimulation (Supplementary Fig. S3A-B). We found that dHepaRG cells maintained without DMSO were able to respond to the action of IFN but not as strongly as dHepaRG maintained in the presence of DMSO.

**Figure 4:**
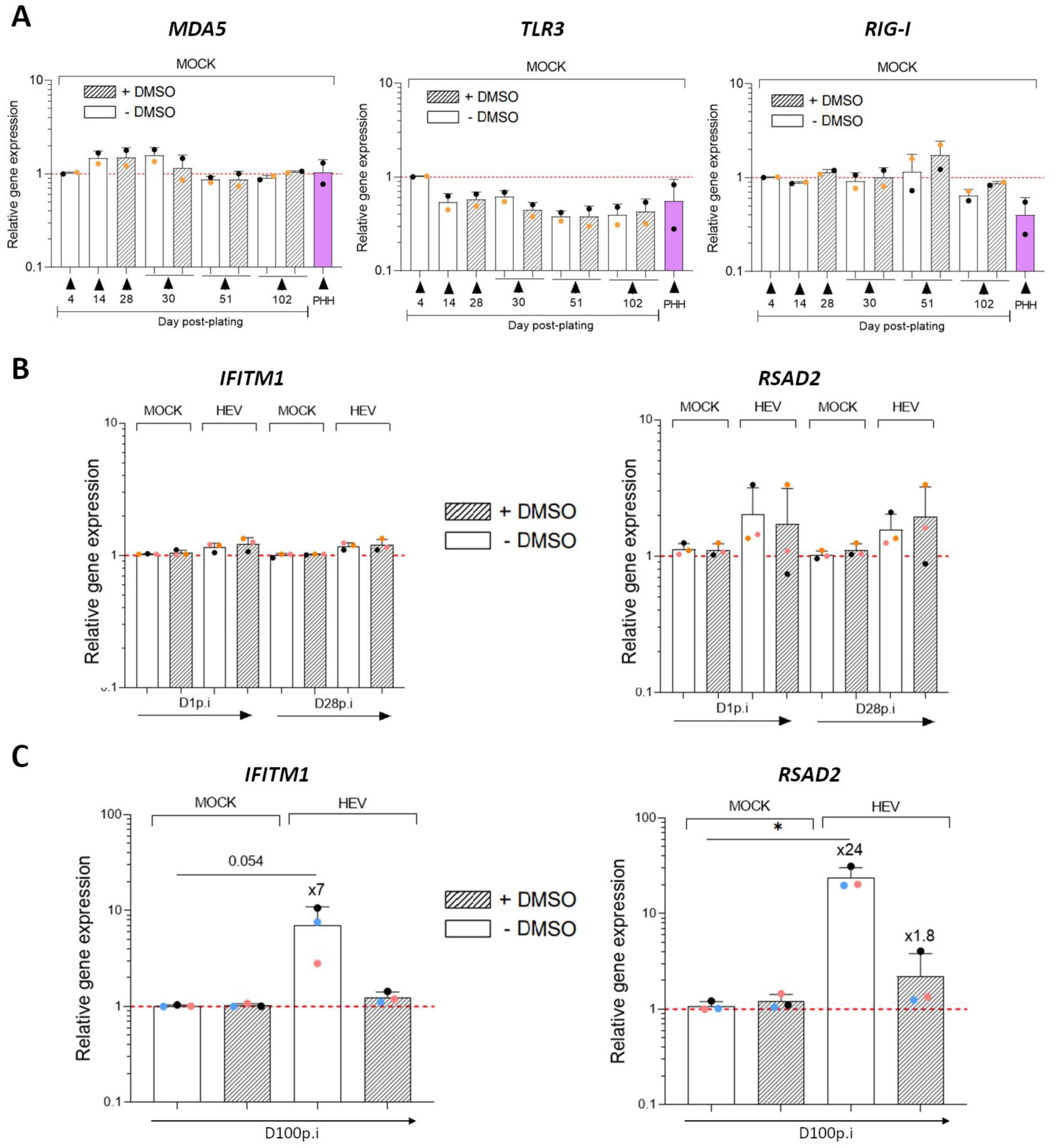
An IFN response is not detected early after HEV-3 infection in dHepaRG. **(A)** Mean relative expression (±SD) of genes coding for innate immune sensors in PHH or in Mock dHepaRG in presence (hatched) or in absence (white) of DMSO upon time. Each independent experiment (n=2) where performed in triplicates. **(B, C)** Mean relative gene expression (±SD) of *IFITM1* and *RSAD2* measured by RT-qPCR in infected (MOI 100) dHepaRG maintained with (hatched) or without (white) DMSO early after infection (D1 and D28p.i) **(B)** or after persistent infection (D100p.i corresponding to D130p.p) **(C)**. Relative gene expression levels were calculated compared to their respective Mock. *GusB (A-C), B2M (A, C)* and *GAPDH (A, C)* were used as housekeeping genes. Each independent experiment (n=3) was performed in triplicates, ANOVA with Dunn’s multiple comparison * *p-value*<0.05, p.p: post-plating, p.i: post-infection.

### 3.6. A stronger IFN response is detected in dHepaRG maintained without DMSO after persistent HEV infection

Next, we wondered whether the presence of DMSO modulate the IFN response in the context of HEV-3 infection. To this end, gene expression of two ISGs was analysed in dHepaRG infected with HEV-3 (MOI 100) and maintained in the presence or not of DMSO for 1, 3, 10 and 28 days p.i (Fig. 4B and Supplementary Fig. S4A). We analysed the expression of *IFITM1* and *RSAD2* as we have previously shown that they are upregulated upon persistent infection in dHepaRG (Meyer et al., 2023). No change in *IFITM1* expression was observed and only a slight upregulation of *RSAD2* expression was detected independently of the presence of DMSO. Intracellular HEV-3 RNA level was, as reported above, higher in cells infected and maintained in absence of DMSO (Supplementary Fig. S4B). Finally, we also measured *IFITM1* and *RSAD2* expression upon persistent infection (D100p.i) in dHepaRG and we observed a significant upregulation of the expression of both ISGs only in cells maintained in absence of DMSO (Fig. 4C). These results suggest that HEV induces low or no innate immune response early after infection in dHepaRG maintained in the presence or absence of DMSO. An IFN response was detected later after infection (D100p.i) only in dHepaRG cultivated in the absence of DMSO when higher level of HEV replication is present (Figure 2B).

## 4. Discussion

Robust and efficient in vitro models supporting HEV replication are needed to improve our knowledge on this hepatotropic virus. Recently, progress have been made to develop pertinent cellular models to study acute and chronic HEV infection such as the HepaRG cell line. Our study provides new insight on the use of this *in vitro* model to study HEV-3 infection. We showed that DMSO-induced differentiation of HepaRG is not essential for HEV-3 infection and that optimal infection is obtained when dHepaRG are infected and maintained in the absence of DMSO during the course of infection. DMSO-induced differentiation and polarization of HepaRG cells did not facilitate HEV infection in HepaRG cells in contrast to what is observed for HBV (Gripon et al., 2002) or human adenovirus infection (Hofmann et al., 2024). This result also differs with several studies showing that DMSO enhances HEV-3 replication in other cell types such as HepG2/C3A and A549 or HEV-C1 and HEV-6 infections in PLC/PRF/5 (Capelli et al., 2020; Chew et al., 2022; Harlow et al., 2024; Primadharsini et al., 2024). This suggests that DMSO impacts HEV infection differently depending on the cell type, culture conditions and/or HEV genotype used. As DMSO modulates the expression of hundreds of genes and acts on many cellular processes such as polarization, antiviral response, metabolism or cell proliferation, it is likely that the chemical modulates some cellular factors that are important for viral entry, egress or replication in a cell-type dependent manner (Dubois-Pot-Schneider et al., 2022; Hofmann et al., 2024; Sugahara et al., 2023).

HepaRG are bipotent cells that can differentiate into hepatocytes and cholangiocytes (Parent et al., 2004). Many studies have described the importance of DMSO in the induction and maintenance of HepaRG differentiation by showing that the chemical compound induces similar expression levels of genes involved in liver metabolism and innate sensing than PHH (Dubois-Pot-Schneider et al., 2022; Gripon et al., 2002; Luangsay et al., 2015). After addition of DMSO, the expression level of multiple hepatocyte-specific genes such as *ALB*, *HNF4α*, *CYP3A4* (Gripon et al., 2002; Noh et al., 2020; Prabhakar et al., 2021) and *G6PC* (Mayati et al., 2018; Wang et al., 2019) is strongly upregulated, as also reported in this study. Commonly used cholangiocyte gene markers are *CK7*, *CK19* and *SOX9* (Dianat et al., 2014; Higuchi et al., 2014; Peng et al., 2018). However, in previous studies, *CK19* expression was shown to be weakly increased (Noh et al., 2020; Prabhakar et al., 2021) and SOX9 expression was decreased (Noh et al., 2020) after DMSO-induced differentiation in HepaRG cells. Other studies have recently shown that these cholangiocyte markers are also present in hepatic-progenitor cells (Aizarani et al., 2019; Lin et al., 2022), suggesting that the interpretation of cholangiocyte markers should be taken with caution. This could explain why, in this study, we have detected no significant change in *CK19* and *SOX9* gene expression in dHepaRG after 2-week exposure to DMSO.

Studies comparing gene expression in presence or in absence of DMSO in dHepaRG have already been performed but only a few days after DMSO removal (Aninat et al., 2006; Dubois-Pot-Schneider et al., 2022). On this short period, they have shown, as reported here, no or only a slight impact of DMSO removal on *ALB* expression and a stronger impact on *CYP3A4* expression (Aninat et al., 2006; Dubois-Pot-Schneider et al., 2022). Nevertheless, the present study shows that expression level of these hepatocyte marker genes stays higher than the one of proliferating HepaRG up to 3.5 months after DMSO removal. Differentiated HepaRG cells cultivated in the absence of DMSO remain then in an intermediate state of differentiation and represent a physiologically pertinent model to study hepatocytes in the context of persistent HEV-3 infection.

MRP2 is one of the canalicular transporter of bile constituents already described as being expressed in polarized HepaRG differentiated with DMSO (Bachour-El Azzi et al., 2015; Le Vee et al., 2013). In our study, we observed by IF a progressive loss of MRP2 in absence of DMSO suggesting a loss of cell polarization after prolonged culture time. Cell polarity has been described as playing an important role in HEV infection. It has been documented that secretion of particles is mainly at the apical side (Le Vee et al., 2013; Sari et al., 2021) with a higher viral infectivity than HEV secreted at the basal side (Capelli et al., 2019; N et al., 2019). However, in 2-dimensional dHepaRG, intra- and extracellular levels of HEV-3 RNA were lower in the presence of DMSO suggesting that cell polarization did not facilitate HEV infection or that its potential enhancement role was hidden by the impact of DMSO on other cellular processes important for HEV infection.

Finally, our study shows that dHepaRG cultivated without DMSO remains a pertinent model to study interaction between HEV and the host innate immune response. We found that non-infected dHepaRG cultivated in the presence or absence of DMSO have similar level of expression of genes coding for intracellular RNA virus sensors and are able to respond to IFN stimulation, even after prolonged time of culture. Independently of the presence of DMSO, we detected no IFN response at early stage of HEV-3 infection. However, upon persistent infection, we detected an increase in the expression of several ISGs only in the absence of DMSO. The absence of IFN response early after infection and in the presence of DMSO could be linked to a lower level of replication and/or percentage of infected cells. Upon HDV infection, dHepaRG are maintained in DMSO and harbor 5 to 10% of infected cells around D5 to D10p.i. This low percentage of infected cells is enough to induce a strong IFN response (Alfaiate et al., 2016; Gillich et al., 2023). This contrasting result between the two hepatic viruses suggests that HEV has evolved strategies to counteract IFN induction and response as already reported in many studies (Bagdassarian et al., 2018; Yadav et al., 2021; Yin et al., 2017).

In conclusion, our study shows that dHepaRG cultivated in the absence of DMSO is a more efficient model to study HEV-3 infection and that this model remains pertinent to study interactions between HEV and the host IFN response over persistent infection. However, more work needs to be done to fully understand the mechanisms behind the detrimental effect of DMSO on HEV infection in dHepaRG, and which stage (s) of the viral cycle is (are) impacted. Such studies could lead to the identification of cellular restriction factors inhibiting HEV replication and the development of better cell culture model for HEV. Such robust cellular models that support long time infection are essential to study HEV-3 infection as this genotype can cause chronic hepatitis.

## Author contributions

Conceptualization, S.G., J.L., D.D., V.D.; Methodology, S.G., M.P., A.MH., J.L., D.D., V.D.; Investigation, S.G., M.P., A.M.H., R.F.; Formal analysis, S.G., V.D.; Resources, M.R., G.P., J.L., D.D., Writing—original draft, S.G., V.D.; Writing—review and editing, S.G., M.P., A.MH., R.F., J.L., D.D., N.P., V.D.; Supervision, V.D. and N.P.; Funding acquisition, V.D, D.D. All authors have read and agreed to the published version of the manuscript.

## Data availability

The data that support the findings of this study are available on request from the corresponding author on reasonable request.

## Acknowledgments

The authors would like to thank Pr Fabien Zoulim, Dr Barbara Testoni and Pr Massimo Levrero for the access to primary human hepatocyte (PHH) isolation platform, as well as Maud Michelet, Anaëlle Dubois, Caroline Pons and Emilie Charles for their help with the isolation of PHH. Moreover, the authors would like to thank Prof. Michel Rivoire’s and Dr. Guillaume Passot’s, and their respective staff in the surgery room, for providing the liver resections. The authors would like to thank also Katja Dinkelborg, Dr. Patrick Behrendt, Dr. George Ssebyatika and Prof. Thomas Krey for their sharing of materials (anti-ORF2 rabbit sera).

