## Supplementary file for "An optimized model for HEV infection in the HepaRG cell line"

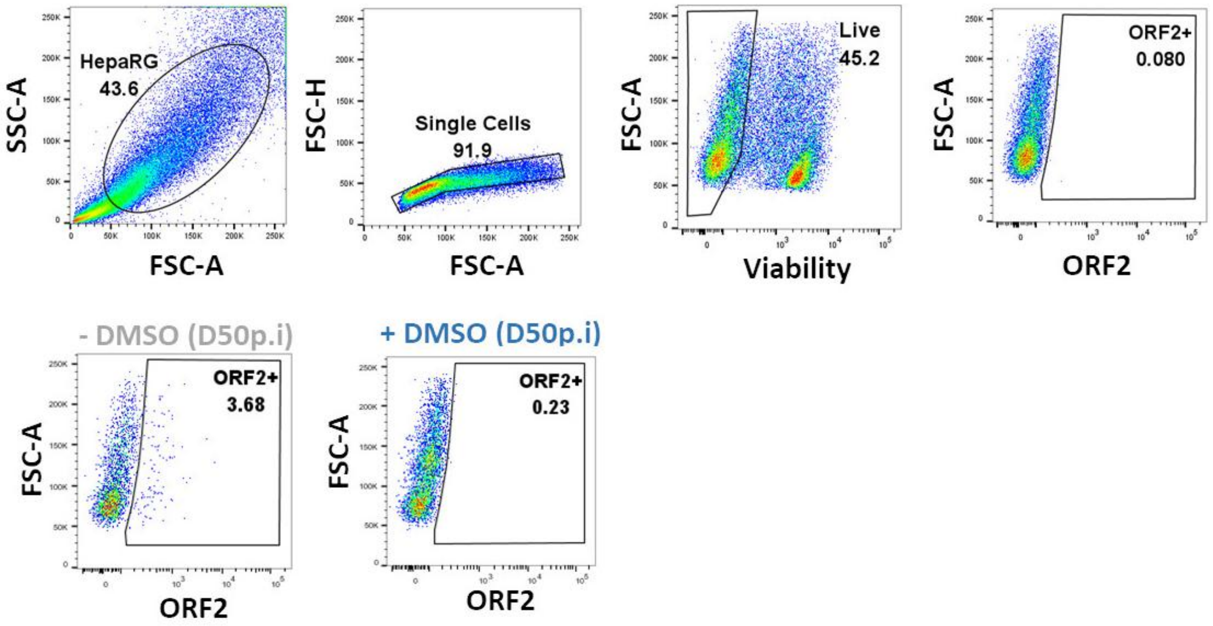


**<u>Supplementary Figure 1:</u>** Percentage of infected (MOI 10, ORF2 staining) dHepaRG maintained in presence (+ DMSO) or in absence of DMSO (- DMSO) measured by flow cytometry after gating (top) in viable cells at D50p.i. p.i: post-infection.

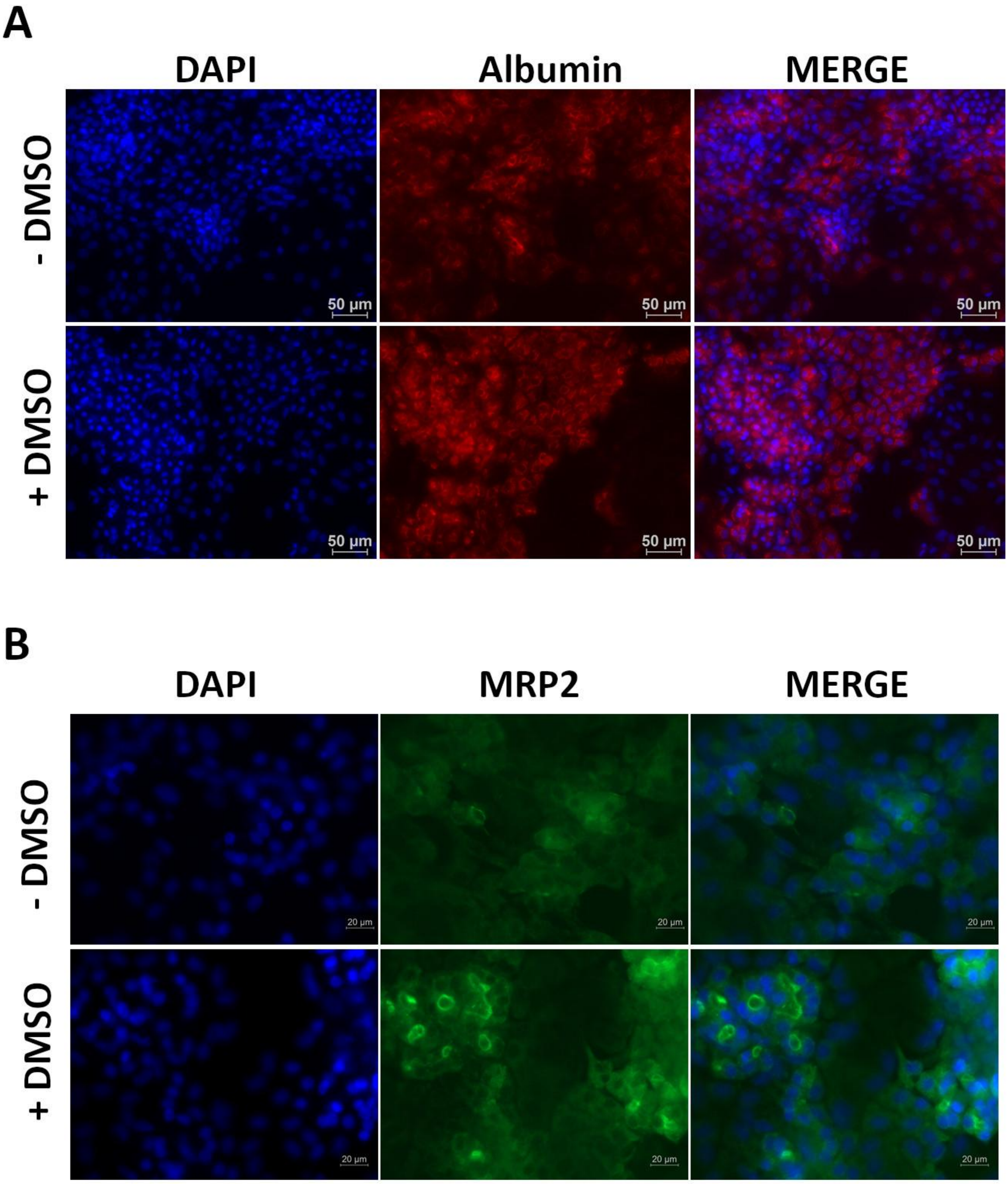


**Supplementary Figure 2: Differentiation and polarization of dHepaRG after DMSO removal**. **(A-B)** Immunofluorescence microscopy on non-infected dHepaRG maintained in presence (+DMSO) or in absence of DMSO (-DMSO). **(A)** Uninfected dHepaRG cells were stained for nuclei (DAPI) and Albumin (hepatocyte marker, red) at D51p.p. Scale bar is 50µm. **(B)** Uninfected dHepaRG cells were stained for nuclei (DAPI) and MRP2 (polarization, green) at D74p.p. Scale bar is 20µm. p.p: post-plating.

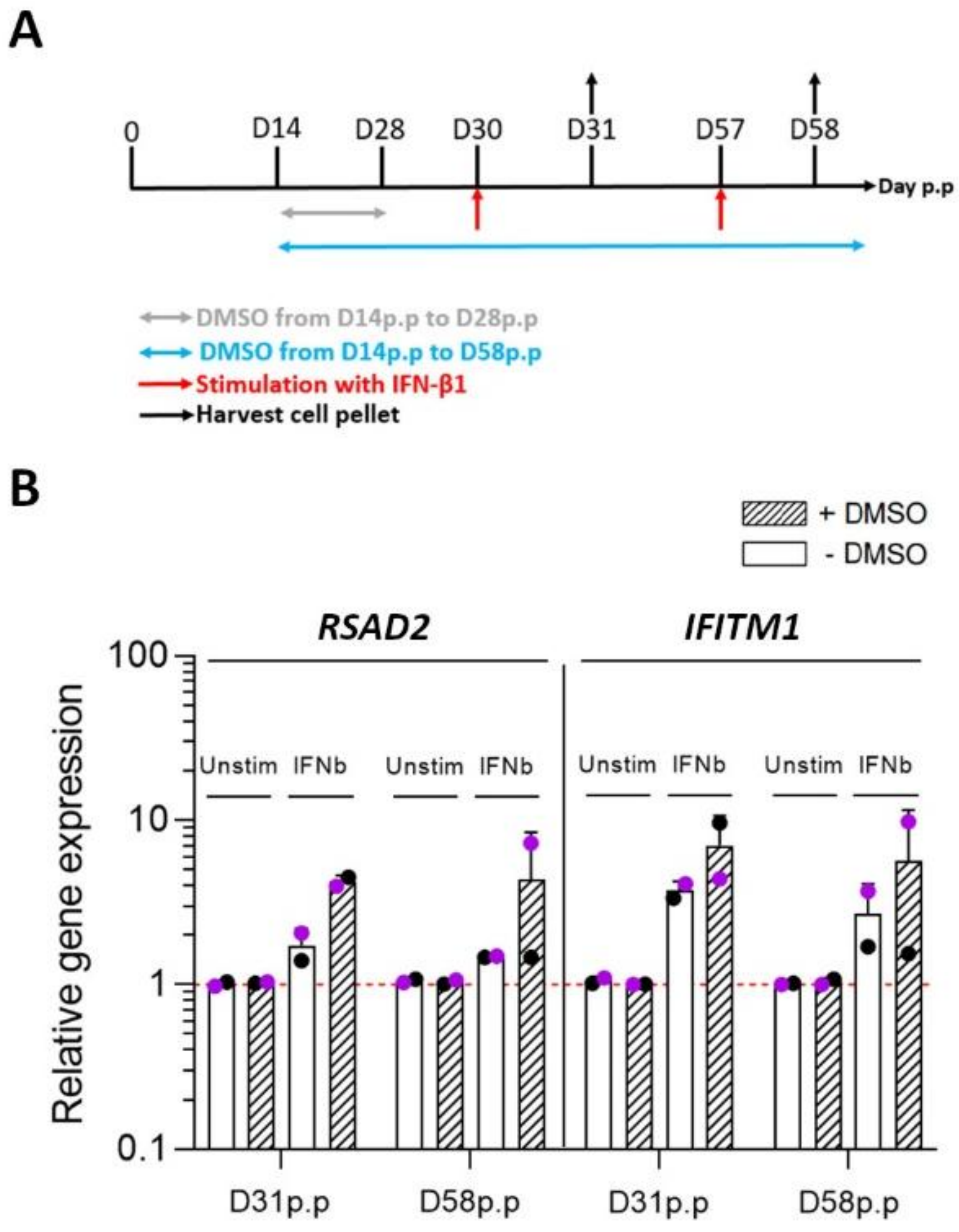


**<u>Supplementary Figure 3: dHepaRG are able to respond to IFN in presence or absence of DMSO.</u> (A)** Uninfected dHepaRG maintained with (hatched) or without (white) DMSO were stimulated overnight with IFN-β1 (200UI/mL) at D30 and D57p.p. The next day, cells were harvested and total intracellular RNA was extracted to quantify ISG mRNA level by RT-qPCR. **(B)** Mean of relative gene expression (±SD) of two ISGs, *IFITM1* and *RSAD2*, compared to their respective unstimulated condition. *GusB* and *B2M* were used as housekeeping genes. Each independent experiment (n=2) was performed in triplicates. p.p: post-plating.

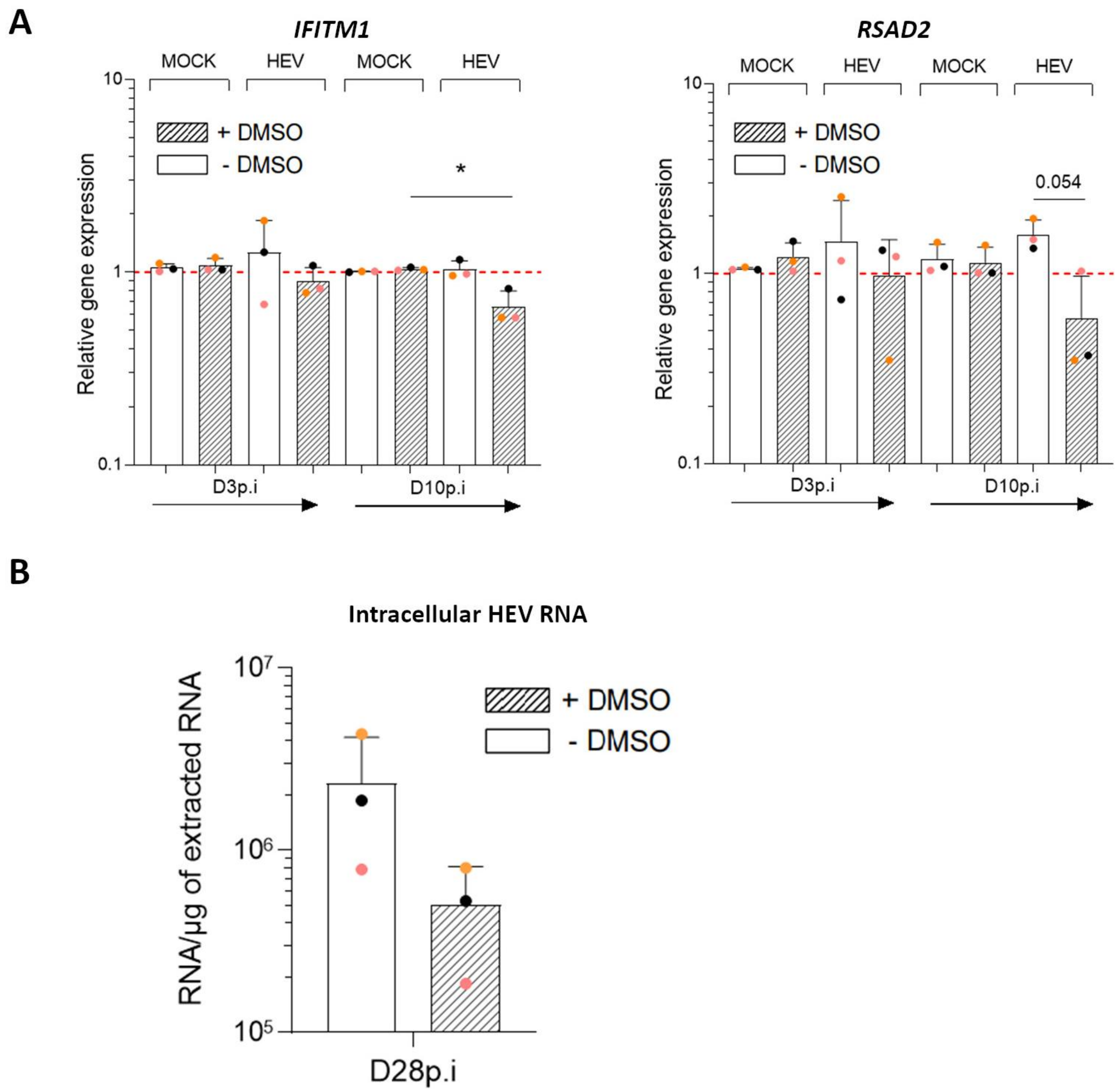


**<u>Supplementary Figure 4:</u> No induction of IFN response early after infection in dHepaRG.** **(A)** Mean of relative gene expression (±SD) of *IFITM1* (left panel) and *RSAD2* (right panel) in infected (MOI 100) dHepaRG at D3 and D10p.i compared to their respective Mock. *GusB* was used as housekeeping gene. Each independent experiment (n=3) was performed in triplicates. ANOVA with Dunn's multiple comparison * *p-value*<0.05. **(B)** Mean of intracellular HEV RNA load (GE/µg of cellular RNA extracted, ±SD) in infected (MOI 100) dHepaRG at D28p.i maintained in presence (hatched) or in absence (white) of DMSO. Each independent experiment (n=3) was performed in triplicates. p.i: post-infection.
